# Impact of AMPK on cervical carcinoma progression and metastasis

**DOI:** 10.1101/2022.07.07.499163

**Authors:** Paweł Konieczny, Tomasz Adamus, Maciej Sulkowski, Marcin Majka

## Abstract

Cervical cancer (CC) is the fourth most common malignant neoplasm among women. Late diagnosis is directly associated with the incidence of metastatic disease and remarkably limits the effectiveness of conventional anticancer therapies at the advanced tumor stage. In this study, we investigated the role of 5’AMP-activated kinase (AMPK) in the metastatic progression of cervical cancer. Since the epithelial mesenchymal transition (EMT) is known as major mechanism enabling cancer cell metastasis, cell lines, which accurately represent this process, have been used as a research model. We used C-4I and HTB-35 cervical cancer cell lines representing distant stages of the disease, in which we genetically modified the expression of the AMPK catalytic subunit α. We have shown that tumor progression leads to metabolic deregulation which results in reduced expression and activity of AMPK. We also demonstrated that AMPK is related to the ability of cells to acquire invasive phenotype and potential for *in vivo* metastases, and its activity may inhibit these processes. Our findings support the hypothesis that AMPK is a promising therapeutic target and modulation of its expression and activity may improve the efficacy of cervical cancer treatment.

## INTRODUCTION

Cervical cancer (CC) is one of the most common malignant neoplasm in women (1). CC mortality affects mainly women in developing countries and is particularly prevalent among patients with a low socio-economic status. An important factor contributing to high mortality is late diagnosis due to poor screening scheme. The main etiological factor of the disease is human papillomavirus (HPV) (2). The development of a vaccine against the most oncogenic types of the virus (16 and 17, which account for 70% of all cases of CC) allows for effective prevention of CC (3). However, vaccination against HPV must be correlated with screening, which allows early detection of pre-cancer lesions. Early stages of the disease are relatively easily curable with conventional therapies such as chemotherapy and radiotherapy, but the chances of 5-year survival dramatically decrease with presence of metastasis (4). Thus, there is unmet need for new therapeutic approaches.

Epithelial to mesenchymal transition (EMT) is long-term morphological and molecular process, allowing cancer cells to lose cell-to-cell adhesion and acquire migratory and invasive properties (5,6). EMT has been demonstrated in pathological conditions such as organ fibrosis and metastatic progression of tumors (7) and it is similar to the physiological process of EMT during embryogenesis, although differences in many types of neoplasms are observed (8).

5’AMP-activated kinase (AMPK) is a cellular energy homeostasis sensor, that coordinates network of various metabolic pathways, such as cholesterol inhibition and fatty acid synthesis (9,10). At molecular level, AMPK controls balance between energy intake and demand, therefore modulating such processes as carbohydrate and lipid metabolism, biosynthesis, autophagy and cell cycle (11). Due to high energy demand during EMT process, several morphological and metabolic changes are orchestrated by AMPK.

Recently, AMPK has emerged as an important therapeutic target for anticancer therapies. Clinical study of type 2 diabetes patients treated with metformin (a pharmacological activator of AMPK), revealed lower cancer incidence rate comparing to the control group (12). Moreover, metformin treatment demonstrated higher cancer remission rate in diabetic patients. (13,14). In vitro studies confirmed the therapeutic potential of AMPK activation in many types of cancers (15,16). Interestingly, AMPK action in cancer progression may be related to blocking or even reversing the EMT.

Current knowledge does not allow unequivocally define the role of AMPK in tumor development and progression, often defining AMPK as a “double-edged sword”, whose action depends on the metabolic context (17,18).

Here, we attempted to elucidate the contribution of AMPK to CC biology, particularly to EMT process and CC metastatic progression.

## RESULTS

### Morphology and growth characteristics of cervical carcinoma cell lines

Morphology and molecular properties of C-4I and HTB-35 cell lines represent different stages of tumor progression - (20). C-4I cells formed colonies with distinct boundaries, and cells were tightly adherent to each other, representing characteristic of epithelial cells, whereas HTB-35 cell line exhibited a loosen cell-to-cell junctions (Fig. 1A). Relative gene quantification showed differences in the expression of major markers associated with the EMT in these cell lines (Fig. 1B). E-cadherin (epithelial cell marker) was significantly decreased, while Vimentin (mesenchymal cell marker) was expressed in abundance in HTB-35 cells. We concluded, that C-4I cell line recapitulates the properties of cells before EMT, while HTB-35 cells that underwent that process.

**Figure 1.**
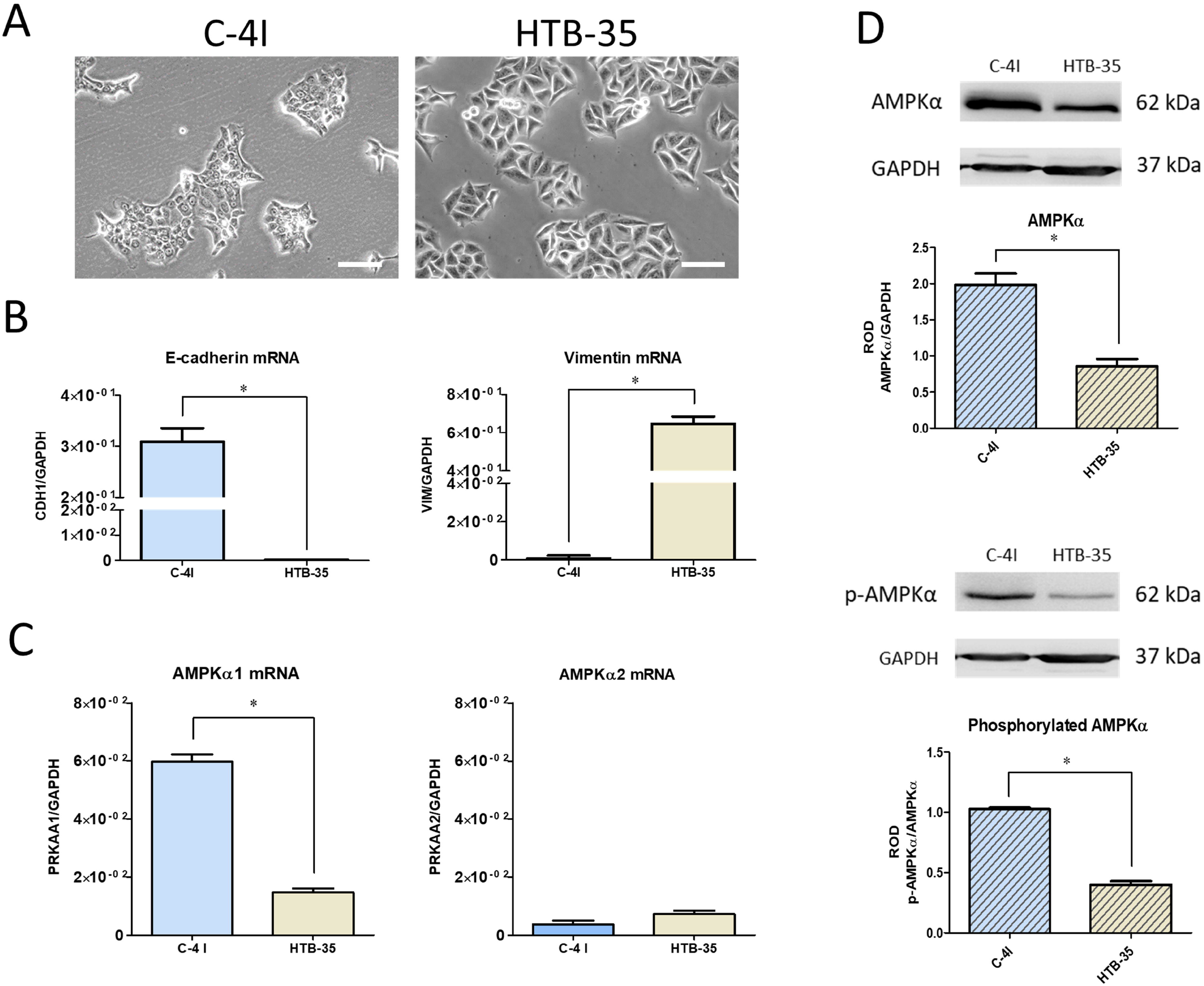
Characteristics of Cervical Carcinoma cell lines. Panel A presents the morphology of CC cells under normal growth conditions (white bar represents 100 μm). Expression of the major epithelial and mesenchymal markers (B) and PRKAAA1 and PRKAA2 genes (C) assessed by qRT-PCR. The graphs represent the mean from at least three independent analyses of the relative expression of the gene to the reference gene GAPDH ± SEM. An asterisk indicates statistically significant differences (p<0.05). Expression and activation of AMPKα protein (D) were evaluated using the Western Blot technique. Membrane pictures show representative results of three independent assessments. Quantitative results presented on bar graphs are based on densitometry of the analyzed gene to the reference gene. Data are presented as mean ± SEM (n=3). Asterisks indicate statistically significant differences (p<0.05).

Next, we assessed AMPK expression level and activity. We showed higher expression level of the AMPKα1 catalytic subunit transcript in the C-4I cell (Fig. 1C). Expression of AMPKα2 subunit transcript was low in both cell lines. Importantly, there were also no differences in the amount of mRNA for β and γ subunits (data not shown). Higher expression of AMPKα in C-4I cells was confirmed by Western Blot (Fig. 1D). Detection of the phosphorylated form of the α subunit confirmed the higher activity of AMPK in C-4I cells (Fig. 1D).

### Effects of EMT inducing factor HGF on AMPK expression and activation

We and others showed previously that hepatocyte growth factor (HGF) is potent EMT inducer in cervical carcinoma (21,22). Thus, we tested the effect of HGF on the expression and activation of the AMPK in CC cells. 24 hours incubation of C-4I cells with HGF induced a mesenchymal phenotype of the C-4I cells when cultured in 10% FBS (Figure 2A). Assuming that the changes of cell shape alternate energy homeostasis, we examined if phenotypic changes were associated with AMPKα expression and activation. Expression of AMPKα1 transcript altered under EMT inducing conditions. The reduction in AMPKα1 expression was statistically significant after 24-hour incubation with HGF, and further increased after 48 hours (Fig. 2B).

**Figure 2.**
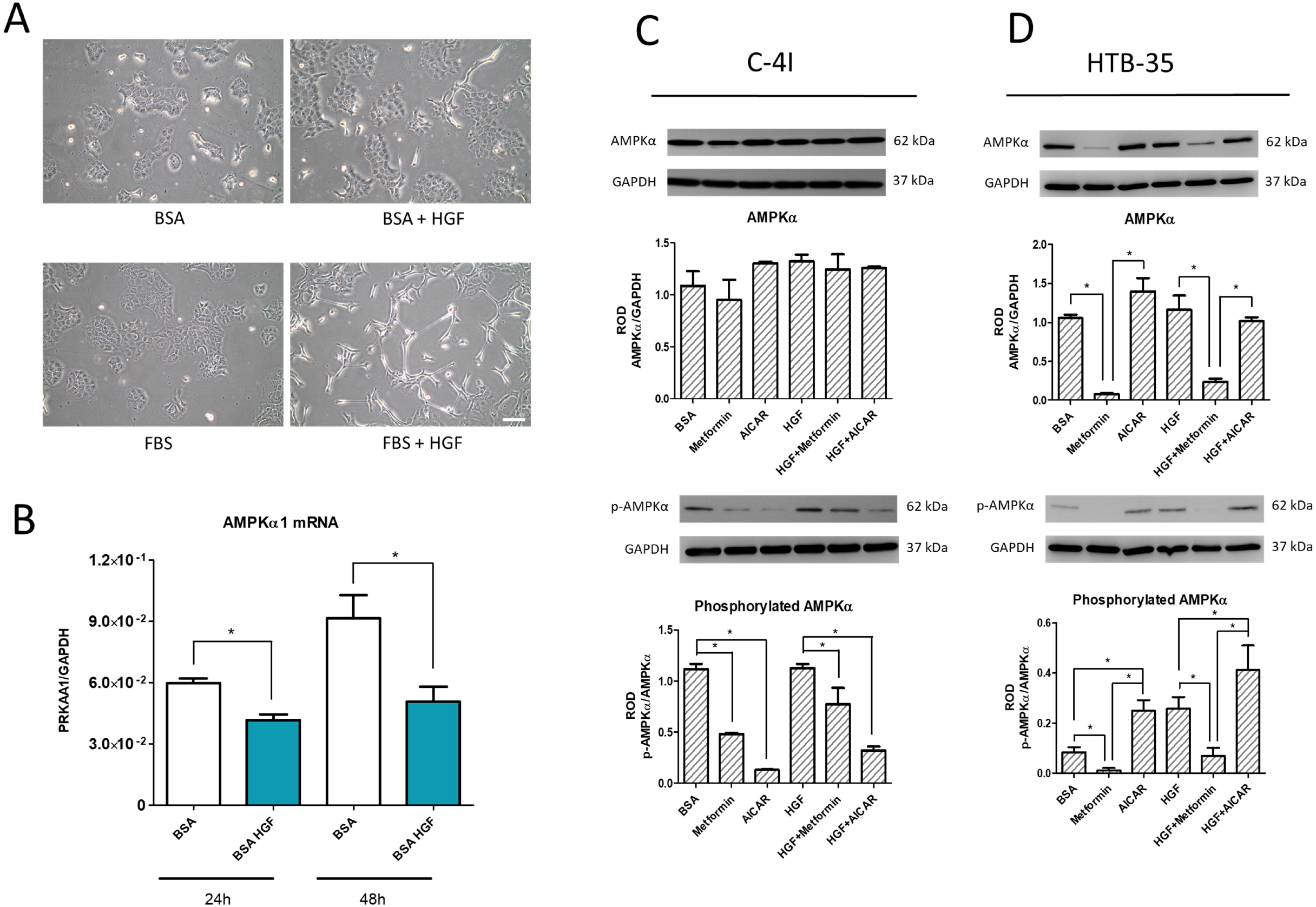
HGF factor influences phenotype of C-4I cells and expression of catalytic subunit AMPKα1. Observation of C-4I cell phenotype change after incubation with HGF, both in medium containing 0.5% BSA and 10% FBS (white bar represents 100 μm). Determination of AMPKα1 transcript level after 24- and 48-hour incubation with HGF in medium containing 0.5% BSA (B). Expression and activation of AMPKα subunit after incubation with HGF and AMPK activators (metformin and AICAR) in C-4I (C) and HTB-35 (D) cells after 24-hour incubation. Membrane pictures show representative results of three independent experiments. Quantitative results presented on bar graphs are based on densitometry of the analyzed gene to the reference gene. Data are presented as mean ± SEM (n=3). Asterisks indicate statistically significant differences (p<0.05).

Following the effect of HGF on the transcript level of the AMPK catalytic subunit α, we evaluated the changes in AMPK protein level after a 24-hour incubation with HGF and pharmacological AMPK activators (Fig. 2C and 2D). We observed, that C-4I cells incubated with metformin and 5-Aminoimidazole-4-carboxamide ribonucleotide (AICAR) for 24-hour lost the activity of the catalytic α subunit. Moreover, this effect was partially reversed by HGF treatment (Fig. 2C).

Surprisingly, we found that C-4I cells simultaneously incubated with HGF and AICAR, did not spread, nor exhibited phenotypic changes (Supplementary Fig. S-1A). Comparable, although weaker effect was observed when C4-I cells were incubated simultaneously with HGF and metformin. For HTB-35 cells, a 24-hour incubation with metformin at a concentration of 10 mM led to partial cell death (Supplementary Fig. S-1C). Interestingly, a very low expression of AMPKα in these cells was found. In addition, virtually no phosphorylated form of AMPKα was detected in cells treated with metformin (Fig. 2D).

### Changes in the expression and activation of the AMPK catalytic subunit a and impact on the expression of markers associated with the EMT

Genetic modifications of the cell lines were used to silence the AMPKα1 subunit in C-4I cells and to upregulate the expression of the AMPKα1 subunit in HTB-35 cell line. To achieve this, we knocked down AMPK subunit α1 expression in C-4I cells using lentiviral particles introducing shPRKAA1 (shAMPKα1). We upregulated expression of AMPKα1 in HTB-35 cells by introducing GFP-P2A-PRKAA1 transgene using lentiviral vectors. We verified downregulation and overexpression of AMPKα1 mRNA level in genetically modified C-4I and HTB-35 cell lines (Fig. 3D) as well as protein level of AMPKα1 (Fig. 3A and 3B). The proliferation of genetically modified CC cells showed no statistically significant changes. Regardless, partial decrease in the growth rate of HTB-35 GFP-P2A-AMPK line cells was observed (Fig. 3C).

**Figure 3.**
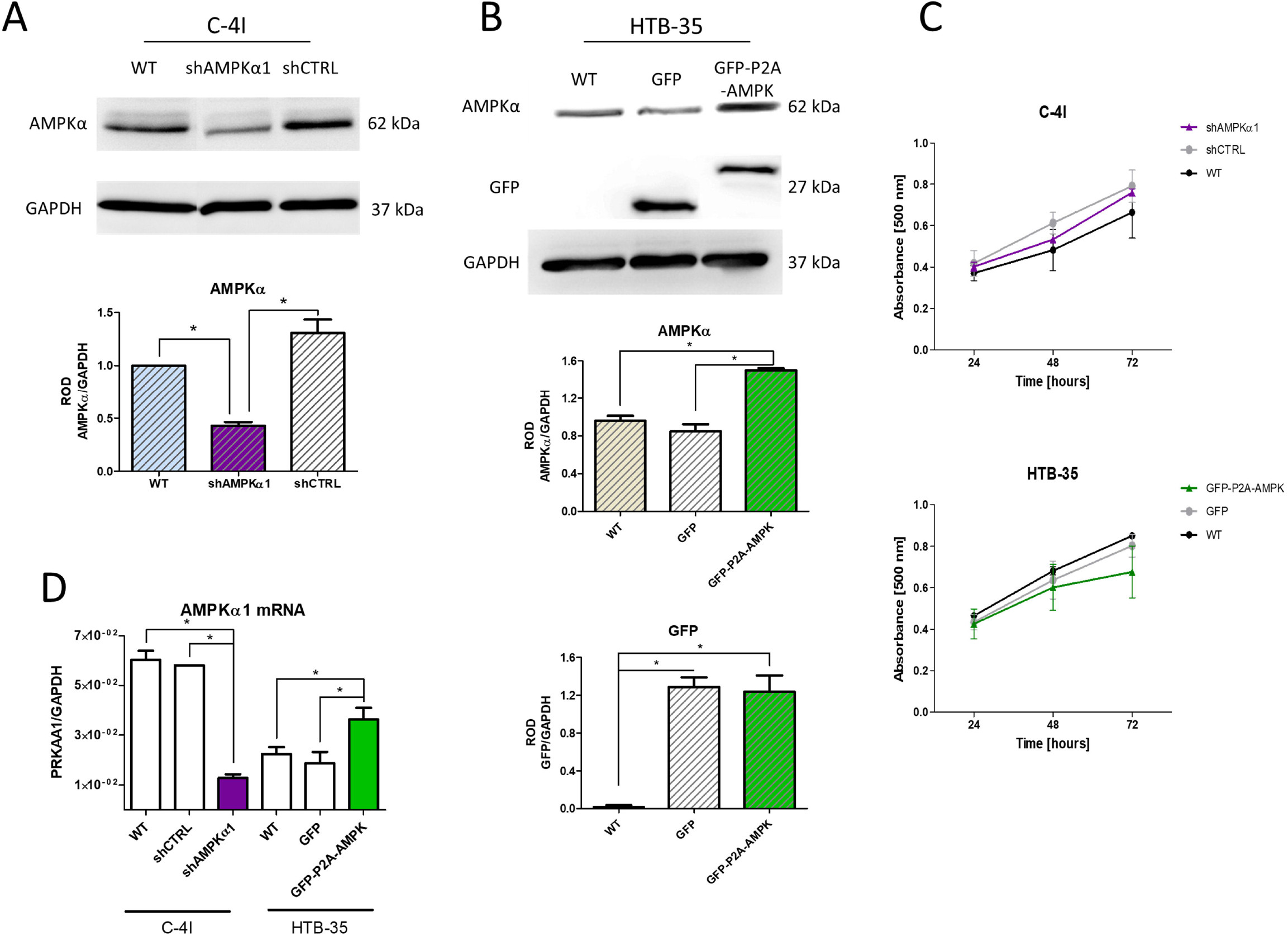
Generation of genetically modified CC lines with increased and decreased AMPKα expression. The evaluation of AMPKα1 catalytic subunit knockdown in C-4I cells (A) and GFP and AMPKα protein expression in selected and sorted HTB-35 lines (B). Membrane pictures show representative results of three independent assessments. Quantitative results presented on bar graphs are based on densitometry of the analyzed gene to the reference gene. Data are presented as mean ± SEM (n=3). Asterisks indicate statistically significant differences (p<0.05). Determination of cell proliferation of modified C-4I and HTB-35 lines by MTS (C). The points on the graph show the mean absorbance of 3 replicates ± SEM. D – transcript level of AMPKα1 catalytic subunit in 6 lines of the developed cellular model. The graphs represent the mean of three independent analyses of the relative expression of the gene to the reference gene GAPDH ± SEM. Asterisks indicate statistically significant differences (p<0.05).

Engineered CC cell lines with modified AMPKα expression served to evaluate the expression of markers associated with EMT. We assessed expression of transcription factors (TEs), as well as changes in mesenchymal phenotype markers or other factors related to EMT process such as intercellular junction proteins.

We discovered significant differences in EMT-related transcription factors in cell lines with altered AMPKα expression. In particular, we observed changes in Snail and ZEB-1 TEs (well-known ‘triggers’ of EMT process) as their expression correlated negatively with AMPKα expression in both C-4I and HTB-35 lines (Fig. 4A and 4C). C4-I shAMPKa1 cells showed elevated levels of Snail and ZEB-1, whereas the opposite effect was observed in the HTB-35 GFP-P2A-AMPK. Elevated AMPKα1 expression in the HTB-35 line correlated with lower expression of the vimentin and higher of E-cadherin, indicating re-establishment of cells epithelial properties (Fig. 4E, 4D). Interestingly, no significant changes in expression of the E-cadherin nor the vimentin were found in C-4I cell lines.

**Figure 4.**
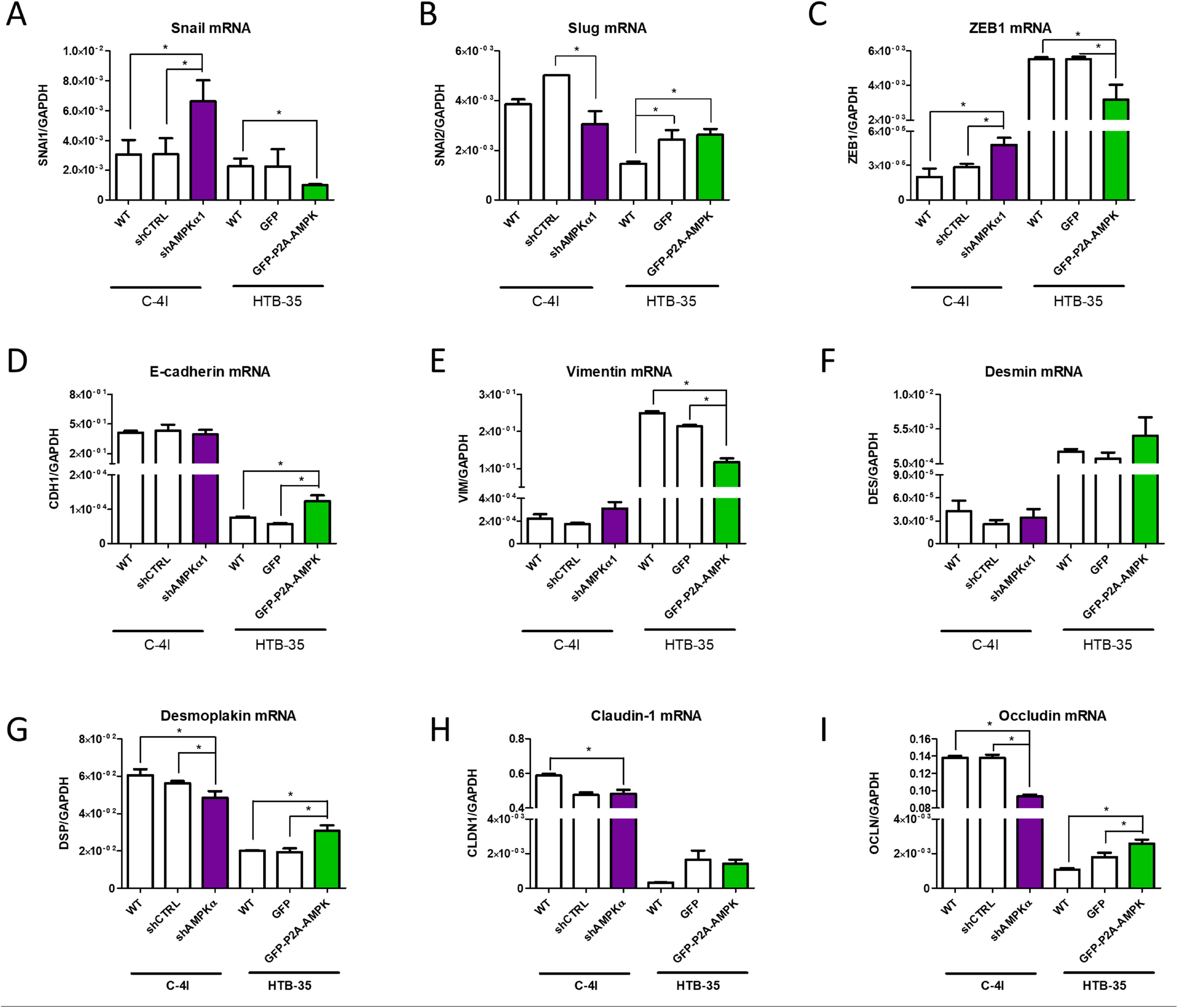
Modification of AMPKα1 catalytic subunit expression in the CC cells alternates the expression of EMT-related factors. Evaluation of expression of transcription factors associated with the induction of epithelial to mesenchymal transition (A, B, C) and transcript level of epithelial and mesenchymal phenotype markers (D, E, F). Transcript level of proteins involved in cell-to-cell connections (G, H, I). The graphs represent the mean of three independent evaluations of the relative expression of the analyzed gene to the reference gene GAPDH ± SEM. Asterisks indicate statistically significant differences (p<0.05).

Our analysis revealed changes in the mRNA expression of the intercellular junction proteins as well. Significant differences in the expression of desmoplakin and occludin were in line with changes of major epithelial and mesenchymal markers (E-cadherin and vimentin). In particular, knockdown of AMPKα decreased the expression of desmoplakin and occludin in C-4I shAMPKα1 cells (Fig. 4G, 4I). Again, the opposite effect was observed for HTB-35 cell lines, as AMPKα overexpression resulted in downregulation of junction proteins.

### Tumor growth and metastatic abilities of C-4I and HTB-35 cell lines

We aimed to verify the influence of AMPKα expression on cells metastatic capacity in NOD-SCID mice. C-4I cells formed tumors when injected subcutaneously into mice (Fig. 5A and Fig. S-4). However, C4-I shAMPK exhibited significantly faster tumor growth than controls, what was corroborated with tumor weight assessed in the end of experiment (Fig. 5B). Despite that, no human β-actin gene transcript was detected in RNA samples collected from mice injected with C-4I cells (Fig. 5C). This indicated the absence of metastasis formation, regardless of AMPK expression.

**Figure 5.**
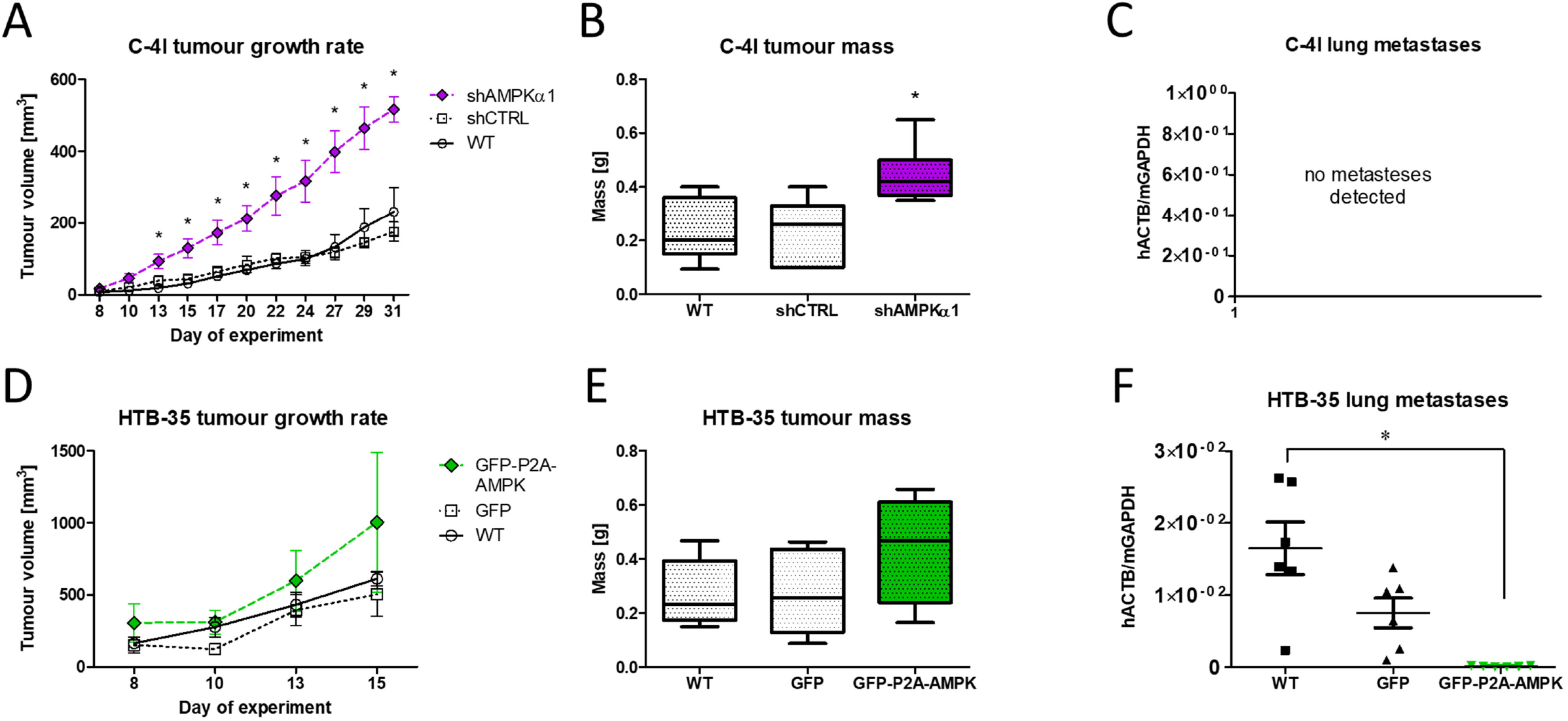
Tumour formation by subcutaneous injection of CC cells in the NOD-SCID mice. The volumetric growth rate of tumours calculated from percutaneous tumour measurements for C-4I (A) and HTB-35 (D) cells. Box plots show the mass of isolated tumors for each of three lines C-4I (B) and HTB-35 (E). Assessment of human β-actin transcript levels in lung samples isolated from mice with C-4I and HTB-35 injected cells (C, F). The ability to form metastases is directly proportional to the amount of human β-actin transcript in relation to the mouse GAPDH gene. All graphs represent data from two rounds of independent experiments ± SEM (2 x n=3 for each cell line). Asterisks indicate statistically significant differences (p<0.05).

We implemented the same experimental approach for HTB-35 cell lines. We found no statistically significant changes in tumor growth and weight (Fig. 5D). The ability of HTB-35 cells to form metastases was assessed by RT-qPCR. Analysis of RNA extracted from lung, spleen and marrow specimens revealed the presence of human β-actin gene transcripts in lungs, indicating HTB-35 cells metastasis. Interestingly, the amount of RNA in the lungs of mice injected with GFP-P2A-AMPK cells was significantly lower in comparison with control groups (Fig. 5F) indicating lower metastatic potential.

## DISCUSSION

The main goal of this study was to correlate the AMPK expression and activation with the acquisition of invasive capabilities through the implementation of EMT.

Our findings revealed that AMPK expression is related to malignant behavior of CC cells. In C-4I line, AMPKα expression and activation stays at a higher level than in HTB-35 cells. Various literature reports show restrained AMPKa expression and activity in multiple types of tumors. Chi-Wai Lee et al. demonstrated a decrease in AMPK expression in different liver cancer cell lines (23). The authors showed that the suppression of AMPKα2 expression in HepG2 cells was associated with an epigenetic modification in the PRKAA2 gene promoter region (encoding AMPKα2). The assessments carried out by Hadad et al. on breast cancer histology samples allowed to correlate the reduced AMPK activity with the development of the tumor (24). In all cases, significantly lower AMPKα activity was determined in tumor tissues than in the surrounding normal epithelial tissue. Additionally, the authors demonstrated that the AMPKα activity is positively correlated with tumor malignant properties. Similar conclusions were reached by researchers analyzing samples of pancreatic cancer tumors (25). IHC staining revealed that AMPKα in cancer cells was deactivated, as opposed to normal epithelium. All above support our hypothesis, that the acquisition of invasive capacity and increase of malignancy rate are positively correlated with the expression and activity of AMPK (especially AMPKα) in CC cells.

Intriguingly, we noticed changes in C4-I cells stimulated with HGF. The induction of EMT with HGF is a well described (21,26,27), but unexpectedly in HGF stimulated C4-I cells, incubation with the AMPK activator AICAR completely blocked scattering effect. Similar but weaker effect was also evident with simultaneous incubation of HGF with either metformin or medium with low glucose concentration, therefore AMPK activation led to inhibition of cell dispersal. To further analyze the influence of HGF on AMPK, the expression and activation of AMPKα at protein level was determined. We found, that in C-4I cells incubated with metformin and AICAR for 24 hours, the activation of AMPKα was diminished. Additional experiments are necessary to fully address this observation. Nevertheless, AMPK activation triggered by HGF remained at a high level but simultaneous incubation with HGF and AMPK activators resulted in lower activation than HGF alone. Remarkably, after 24 hours of incubation in the medium with 0.5% BSA (without any additional factors added), AMPKα appeared to be activated due to induced nutrient deficiency in the medium and natural activation (28).

We could explain observed phenomena by the fact that C-4I cells, as retaining some features of normal epithelial cells, also maintain a metabolic ‘regime’. This means that long-term and non-physiological activation of the AMPK will be unfavorable for them because the AMPK loses its proper function, that is the metabolic switch - continuous activation makes it impossible to react to changes. Therefore, the cells switch off the AMPK activity. However, the influence of HGF bypasses this mechanism and leads to permanent activation of AMPKα, probably by changing the concentration of Ca2+ ions and activation of CaMKK (29,30). Hence, incubation with HGF (in a medium with 10% FBS) leads to the re-arrangement of the cytoskeleton and the acquisition of invasive phenotype, as energy is still supplied by AMPK. In turn, the observed silencing of AMPK activity in case of simultaneous incubation with AICAR and HGF explains the inhibition of cell dispersion by switching the AMPK from energy generation to energy storage processes.

AMPK activation by HGF and pharmacological AMPK activators in HTB-35 cells appears to be completely different than in C-4I cells. It is possible that we observed an active mechanism degrading the catalytic subunit of AMPKα. Studies conducted by Pineda et al. shows that MAGE-A3/6-TRIM28 ubiquitous ligase caused ubiquitination and subsequent degradation of α1 subunit in different types of neoplasms (31). Importantly, it was associated with the hypersensitivity of cancer cells to AMPK activator - metformin. The occurrence of this mechanism in HTB-35 cells could explain their high sensitivity to long-term incubation with metformin. We also observed that effect for AICAR stimulated cells, however it was visible after 48 to 72 hours of incubation.

We illustrated dual nature of AMPK in examined CC cells, most likely resulting from differences in cell invasiveness and malignancy (32). HTB-35 cells utilize AMPK not as a “guardian” of metabolism, but as a “helper” in gaining energy, e.g., through disorders in lipid metabolism. However, the use of metformin, whose action is less selective than AICAR, in HTB-35 cells leads to a decrease in the efficiency of energy acquisition in the process of glycolysis and, consequently, cell death (33).

We noticed that different levels of AMPK expression leads to different expression levels of intercellular junction proteins. RNA expression for desmoplakin and occludin was elevated in HTB-35 cells with AMPKα overexpression. Noteworthy, AMPKα knockdown in C-4I cells resulted in exactly opposite effect – lowered transcript level of intercellular junction proteins was detected. These observations underline AMPK impact on phenotype change during EMT. Moreover, we conclude that the reduction of AMPK expression is correlated with the first step for acquisition of motility properties. The changes in AMPK expression have also an impact on transcription factors associated with EMT. AMPKα1 knockdown in C-4I cells increased the expression of Snail and ZEB-1 factors, while increased AMPKα1 expression in HTB-35 cells led to decrease levels of these TFs. Moreover, HTB-35 cell line overexpressing AMPKα shows differences in the expression of the main EMT markers such as vimentin and E-cadherin, showing a partial reversal of mesenchymal phenotype. C-4I shAMPKα1 cell line, apart from changes in the expression of transcription factors, did not show significant differences in E-cadherin or vimentin. This suggests that the decrease in AMPK expression alone is not sufficient to trigger changes in the EMT phenotype. This is consistent with the literature reports, connecting the reduction of this kinase activity with the acquisition of invasive capacity by cancer cells (34).

Although we did not notice significant changes in the growth rate between unmodified and modified cells in vitro, AMPKα knockdown in C-4I cells enhanced their ability to grow in vivo. After transplantation to the NOD-SCID mouse model, C4-I shAMPKa1 cells formed larger and faster growing tumors than control cells. This is consistent with the published data, which correlates the expression of AMPKα directly with the tumor growth rate in the mouse model (35). However, the silencing of AMPKα expression did not affect the lack of ability of C-4I cells to metastasize. Considering the lack of significant changes of EMT major markers (E-cadherin and vimentin) in these cells, and the fact that EMT is the necessary mechanism for the metastasis in CC (36), this result is not surprising.

Conversely, an increase of AMPKα1 expression in HTB-35 cells, which was associated with a partial reversal of the mesenchymal phenotype in vitro, limited the ability of these cells to metastasize in vivo. Considering the fact that this kinase is natively active in HTB-35 cells (although at a low level), the increase in the amount of AMPKα protein will simultaneously result in proportional increase in its activity. It is in accordance with most of the publications, indicating that of AMPK activation leads to a decrease metastatic capability (25,37).

AMPK can take on different functions depending on the molecular and metabolic context (38,39). We showed here that lowering the expression of AMPKα in C-4I cells and overexpression in HTB-35 cells lead to the promotion of cells survival in vivo. This may be due to the different function of AMPK in these cells. In C-4I line, which retains some of the properties of normal epithelial tissue, AMPK still plays the correct role of metabolic “guardian”, whose exclusion leads to metabolic deregulation and acquisition of a higher degree of malignancy. In HTB-35 cells, there is a permanent decoupling of metabolic pathways, therefore AMPK falls out of its function. HTB-35 cells, which acquire energy mainly in the process of glycolysis (40), can use AMPK only to increase the ability to generate energy, without metabolic regulation (41).

Dissecting the role of AMPK in regulating many intracellular processes is still a demanding issue. Despite the fact that AMPK may play a double role in the cancer progression, it is a promising therapeutic target. Especially in the early stages of CC, the AMPK activators may benefit in favorable therapeutic outcome. In the advanced neoplastic disease, or to be more precise, in the case of deregulation the expression and activation of AMPK in CC cells, the restoration of AMPK expression may be a step towards the elimination of cancer cells.

## MATERIAL AND METHODS

### Cell culture conditions

C-4I and HTB-35 cell lines were maintained as a monolayer culture in Waymouth’s medium (Thermo Fisher Scientific, MA, USA) and EMEM medium (Lonza, Switzerland) respectively, supplemented with 10% v/v fetal bovine serum (FBS; EURx, Poland), and 100 U/ml penicillin and 100 μg/ml streptomycin (both from Thermo Fisher Scientific, Waltham, MA, USA). Cells were cultured at 37°C in a humidified atmosphere of 5% CO_2_.

### Genetic modification of C-4I and HTB-35 cells

Cloning and lentivirus production was previously described in (19). Further details of modified cell lines generation are given in **Supplementary Materials.**

### qRT-PCR

Total RNA was isolated with GeneMATRIX Universal RNA kit (Eurx, Poland), followed by reverse transcription (RT) using M-MLV reverse transcription kit (Promega, Madison, WI, USA). Blank qPCR Master Mix kit (EURx, Poland) with specific TaqMan probes (Thermo Fisher Scientific, USA) were used (*Table 1*). The ddCt method (2^-ΔΔCt^) was used to calculate relative expression of the genes.

### Western Blot analysis

The total protein fraction was extracted using M-PER buffer (Thermo Fisher Scientific, USA) with protease and phosphatase inhibitors (Sigma Aldrich, USA). The protein concentration was determined by Bradford method. After SDS-PAGE and proteins transferred on PVDF membranes were incubated overnight at 4°C with primary antibodies (*Table 2*), and subsequently detected with HRP-conjugated goat anti-rabbit IgG secondary antibody (1:4000; Santa Cruz Biotechnology, USA). The membranes were developed with SuperSignal West Pico Chemiluminescence Substrate (Thermo Fisher Scientific, USA) with Gel Logic Imaging System (Kodak, USA).

### MTS proliferation assay

The CellTiter 96^®^ AQueous One Solution Cell Proliferation Assay kit (Promega, USA) was used to assess the proliferation of the CC cells. Cells were seeded into 96-well plates - the number of cells seeded was 1×10^4^ and 5×10^3^ for C-4I and HTB-35 cell lines respectively. The assay was performed at three time points: 24 hours, 48 hours, and 72 hours after cell seeding.

### Xenografts in NOD-SCID mouse model

Animal experiments were conducted in accordance to ethical committee guidelines with approval of Local Institutional Animal Care and Use Committee (IACUC) in Krakow. 2×10^6^ C-4I cells(WT, shCTRL and shAMPKα1) or 1×10^6^ HTB-35 (WT, GFP,GFP-P2A-AMPK) were injected in 200μL of PBS/growth factor-reduced Matrigel (Corning) 1:1 into left dorsal flank of adult female NOD/SCID mice. When tumor became palpable, its size was measured every 2-3 days. Experiments were terminated when tumor volume exceeded 1000 mm^3^ or earlier based on animal welfare. Mice were sacrificed, tumors, bone marrow, spleen, liver, and lungs were collected and froze in LN2. Prior to RNA extraction, the frozen tissue fragments were mechanically homogenized in a Tissuelyser II device (Qiagen, Germany) and total RNA was isolated with GeneMATRIX Universal RNA Purification Kit (EURx, Poland).

### Statistical analysis

Statistical analysis was performed using GraphPad Prism 5. If not stated otherwise, presented data represents 3 independent experiments. For comparisons of 2 groups of data, the Student t test was used. For comparisons of multiple groups, analysis of variance ANOVA with Tukey’s post-hoc test was used. p=0.05

## Acknowledgements

We would like to acknowledge Kazimierz Weglarczyk and Rafal Szatanek for technical help with BD FACSAria; Marta Kot for help with Attune Flow Cytometer; Klaudia Skrzypek for revising the manuscript. The project was supported by the research grants from Jagiellonian University Medical College to MM: K/ZDS/003725

## Author contributions

P.K. designed, planned and conducted most of the experiments *in vitro* and *in vivo*, analyzed data, performed the statistical analysis and wrote the manuscript. T.A. was involved in molecular cloning of GFP-P2A-PRKAA, generation of plasmids and viral vectors. M.S. was involved in viral vectors generation and *in vivo* experiments. M.M. conceived the study, designed and coordinated the study, wrote manuscript. All the authors revised the manuscript.

## Conflict of interest

The authors declare that they have no conflict of interest.

## Notes

### Competing Interest Statement

The authors have declared no competing interest.

## REFERENCES

1. Wild CP, Weiderpass E, Stewart BW. World Cancer Report: Cancer Research for Cancer Prevention. 2020

2. Bosch FX, De Sanjosé S. The epidemiology of human papillomavirus infection and cervical cancer. Dis. Markers. 2007; 23: 213–227.

3. Tran NP, Hung C-F, Roden R, Wu T-C, Tran NP, Hung C-F et al. Control of HPV Infection and Related Cancer Through Vaccination. Recent Results Cancer Res 2014; 193:149–171.

4. Cohen PA, Jhingran A, Oaknin A, Denny L. Cervical cancer. Lancet (London, England) 2019; 393: 169–182.

5. Kalluri R. EMT: When epithelial cells decide to become mesenchymal-like cells. J Clin Invest 2009; 119: 1417–1419.

6. Lamouille S, Xu J, Derynck R. Molecular mechanisms of epithelial-mesenchymal transition. Nat. Rev. Mol. Cell Biol. 2014; 15: 178–196.

7. Thiery JP, Acloque H, Huang RYJ, Nieto MA. Epithelial-Mesenchymal Transitions in Development and Disease. Cell. 2009; 139: 871–890.

8. Nisticò P, Bissell MJ, Radisky DC. Epithelial-mesenchymal transition: General principles and pathological relevance with special emphasis on the role of matrix metalloproteinases. Cold Spring Harb Perspect Biol 2012; 4.

9. Oakhill JS, Scott JW, Kemp BE. Structure and function of AMP-activated protein kinase. Acta Physiol 2009; 196: 3–14.

10. Sanz P. AMP-activated protein kinase: structure and regulation. Curr Protein Pept Sci 2008; 9: 478–92.

11. Tyszka-Czochara M, Konieczny P, Majka M. Recent advances in the role of AMP-activated protein kinase in metabolic reprogramming of metastatic cancer cells: targeting cellular bioenergetics and biosynthetic pathways for anti-tumor treatment. J. Physiol. Pharmacol. 2018; 69.

12. Evans JMM, Donnelly LA, Emslie-Smith AM, Alessi DR, Morris AD. Metformin and reduced risk of cancer in diabetic patients. Br Med J 2005; 330: 1304–1305.

13. Coyle C, Cafferty FH, Vale C, Langley RE. Metformin as an adjuvant treatment for cancer: a systematic review and meta-analysis. Ann Oncol Off J Eur Soc Med Oncol 2016; 27: 2184–2195.

14. Zhong S, Wu Y, Yan X, Tang J, Zhao J. Metformin use and survival of lung cancer patients: Meta-analysis findings. Indian J Cancer 2017; 54: 63–67.

15. Fogarty S, Hardie DG. Development of protein kinase activators: AMPK as a target in metabolic disorders and cancer. Biochim Biophys Acta - Proteins Proteomics 2010; 1804: 581–591.

16. Luo Z, Zang M, Guo W. AMPK as a metabolic tumor suppressor: Control of metabolism and cell growth. Futur. Oncol. 2010; 6: 457–470.

17. Jeon SM, Hay N. The double-edged sword of AMPK signaling in cancer and its therapeutic implications. Arch Pharm Res 2015; 38: 346–357.

18. Zadra G, Batista JL, Loda M. Dissecting the dual role of AMPK in cancer: From experimental to human studies. Mol. Cancer Res. 2015; 13: 1059–1072.

19. Adamus T, Konieczny P, Sekuła M, Sułkowski M, Majka M. The strategy of fusion genes construction determines efficient expression of introduced transcription factors. Acta Biochim Pol 2014; 61.

20. Fogh J, Giovanella B. The Nude Mouse in Experimental and Clinical Research. Jorgen Fogh, Beppino C. Giovanella. Q Rev Biol 1979; 54: 96–96.

21. Miekus K, Pawlowska M, Sekuła M, Drabik G, Madeja Z, Adamek D et al. MET receptor is a potential therapeutic target in high grade cervical cancer. Oncotarget 2015; 6:10086–10101.

22. Boromand N, Hasanzadeh M, ShahidSales S, Farazestanian M, Gharib M, Fiuji H et al. Clinical and prognostic value of the C-Met/HGF signaling pathway in cervical cancer. J Cell Physiol 2018; 233: 4490–4496.

23. Lee CW, Wong LLY, Tse EYT, Liu HF, Leong VYL, Lee JMF et al. AMPK promotes p53 acetylation via phosphorylation and inactivation of SIRT1 in liver cancer cells. Cancer Res 2012; 72: 4394–4404.

24. Hadad SM, Baker L, Quinlan PR, Robertson KE, Bray SE, Thomson G et al. Histological evaluation of AMPK signalling in primary breast cancer. BMC Cancer 2009; 9: 307.

25. Chen K, Qian W, Li J, Jiang Z, Cheng L, Yan B et al. Loss of AMPK activation promotes the invasion and metastasis of pancreatic cancer through an HSF1-dependent pathway. Mol Oncol 2017; 11: 1475–1492.

26. Farrell J, Kelly C, Rauch J, Kida K, García-Muñoz A, Monsefi N et al. HGF Induces Epithelial-to-Mesenchymal Transition by Modulating the Mammalian Hippo/MST2 and ISG15 Pathways. J Proteome Res 2014; 13: 2874–2886.

27. Liu F, Song S, Yi Z, Zhang M, Li J, Yang F et al. HGF induces EMT in non-small-cell lung cancer through the hBVR pathway. Eur J Pharmacol 2017; 811: 180–190.

28. Hardie DG, Ross FA, Hawley SA. AMPK - a nutrient and energy sensor that maintains energy homeostasis. Nat Rev Mol Cell Biol 2012; 13: 251.

29. Majka M, Drukala J, Lesko E, Wysoczynski M, Jenson AB, Ratajezak MZ. SDF-1 alone and in co-operation with HGF regulates biology of human cervical carcinoma cells. Folia Histochem Cytobiol 2006; 44: 155–164.

30. Chou CC, Lee KH, Lai IL, Wang D, Mo X, Kulp SK et al. AMPK reverses the mesenchymal phenotype of cancer cells by targeting the Akt-MDM2-Foxo3a signaling axis. Cancer Res 2014; 74: 4783–4795.

31. Pineda CT, Ramanathan S, Fon Tacer K, Weon JL, Potts MB, Ou YH et al. Degradation of AMPK by a cancer-specific ubiquitin ligase. Cell 2015; 160: 715–728.

32. Tyszka-Czochara M, Lasota M, Majka M. Caffeic acid and metformin inhibit invasive phenotype induced by TGF-β1 in C-4I and HTB-35/SiHa human cervical squamous carcinoma cells by acting on different molecular targets. Int J Mol Sci 2018; 19.

33. Tyszka-Czochara M, Bukowska-Strakova K, Majka M. Metformin and caffeic acid regulate metabolic reprogramming in human cervical carcinoma SiHa/HTB-35 cells and augment anticancer activity of Cisplatin via cell cycle regulation. Food Chem Toxicol 2017; 106: 260–272.

34. Li W, Saud SM, Young MR, Chen G, Hua B. Targeting AMPK for cancer prevention and treatment. Oncotarget. 2015; 6: 7365–7378.

35. Cheng J, Huang T, Li Y, Guo Y, Zhu Y, Wang Q et al. AMP-activated protein kinase suppresses the in vitro and in vivo proliferation of hepatocellular carcinoma. PLoS One 2014; 9: 1–10.

36. Lee MY, Chou CY, Tang MJ, Shen MR. Epithelial-mesenchymal transition in cervical cancer: Correlation with tumor progression, epidermal growth factor receptor overexpression, and snail up-regulation. Clin Cancer Res 2008; 14: 4743–4750.

37. Cao W, Li J, Hao Q, Vadgama J V, Wu Y. AMP-activated protein kinase: A potential therapeutic target for triple-negative breast cancer 11 Medical and Health Sciences 1112 Oncology and Carcinogenesis. Breast Cancer Res. 2019; 21: 29.

38. Chuang H-C, Chou C-C, Kulp SK, Chen C-S. AMPK as a potential anticancer target - friend or foe? Curr Pharm Des 2014; 20: 2607–18.

39. Faubert B, Vincent EE, Poffenberger MC, Jones RG. The AMP-activated protein kinase (AMPK) and cancer: Many faces of a metabolic regulator. Cancer Lett 2015; 356:165–170.

40. Tyszka-Czochara M, Bukowska-Strakova K, Kocemba-Pilarczyk KA, Majka M. Caffeic acid targets AMPK signaling and regulates tricarboxylic acid cycle anaplerosis while metformin downregulates HIF-1α-induced glycolytic enzymes in human cervical squamous cell carcinoma lines. Nutrients 2018; 10: 841.

41. Vara-Ciruelos D, Russell FM, Hardie DG. The strange case of AMPK and cancer: Dr Jekyll or Mr Hyde? Open Biol 2019; 9: 190099.

