## Supplementary materials for "Impact of AMPK on cervical carcinoma progression and metastasis"

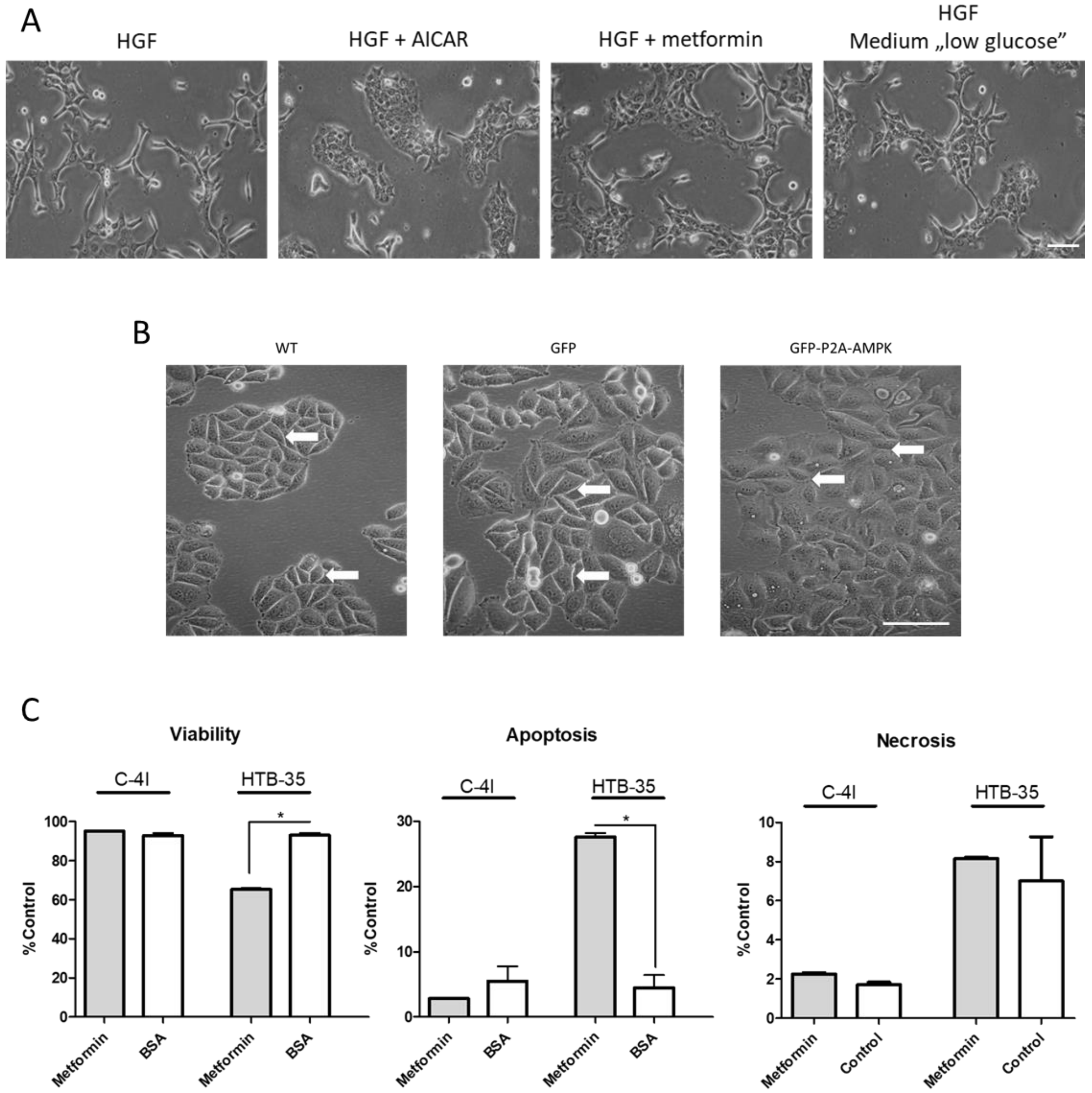


**Figure S- 1.** Differences in colony apperance in C-4I cells stimulated with HGF with and without AMPK activators [medium containing 10% FBS] (A). Chganges in cell-to-cell conections in HTB-35 GFP-P2A-AMPK cell line. Visible differences marked with white arrows (B). White bars represent 100 µm. Metformin stimaulation of C-4IWT and HTB-35 WT resulted in viability decrease, apoptosis and necrosis induction for HTB-35 WT cells. Data are presented as mean ± SEM (n=3). Asterisks indicate statistically significant differences ($p<0.05$).

### *C-4I modification*

To obtain modified C-4I cell lines, cells were infected using purchased lentiviruses introducing shRNAs against the catalytic subunit of AMPKα1 and, as a control, shRNAs not targeting any known transcript ("scrambled" shRNAs). Transduction was performed using MOI=5 (Multiplicity of infection) and 6 µg/ml polybrene (Sigma-Aldrich). Selection of C-4I modified shCTRL and shAMPKα1 cells was performed using the selection antibiotic puromycin (Invivogen, USA) at a concentration of 0.125 µg/ml. The appropriate concentration of antibiotic was chosen based on the survival curve performed previously on the cells of this line. Antibiotic selection of C-4I shCTRL and shAMPKα1is illustrated in Figure S-2.

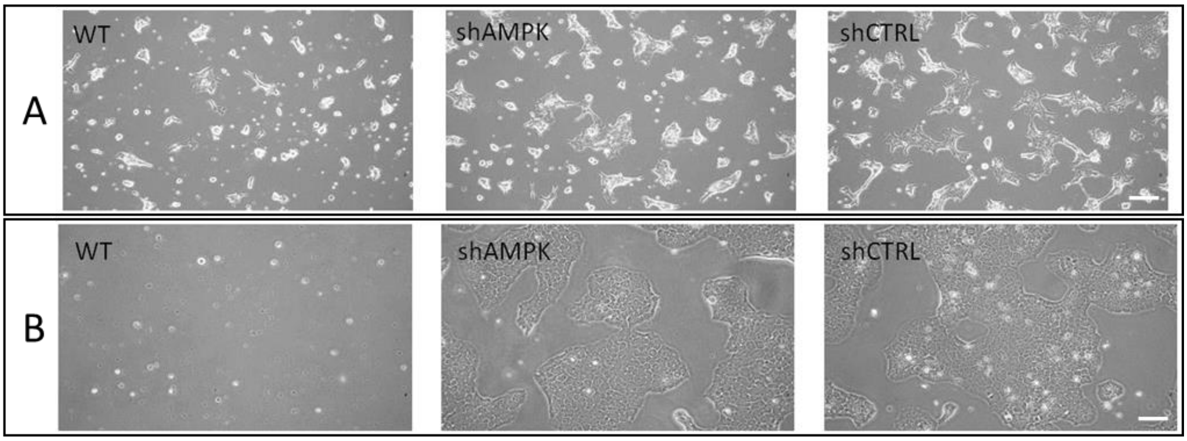


**Figure S- 2.** Antibiotic selection of C-4I shAMPK and shCTRL cell lines. Panel A – 5 days of puromycin selection. Panel B – 10 days of puromycin selection – no WT cells visible (end of selection). White bars represent 100 µm.

### *Creation of the GFP-P2A-AMPKα1 gene construct*

The GFP-P2A-AMPKα1 gene construct was obtained, by PCR reactions introducing both restriction sites and recombination sequences (attB) at the insert ends. The green fluorescent protein (GFP) sequence was amplified from the pEGFP-C1 plasmid (Clonetech, USA), the sequence of the α1 catalytic subunit of AMPK kinase from the pDONR223-PRKAA1 [#23871] plasmid (Addgene, USA), and P2A from the FUW-OSKM plasmid (Addgene, USA). Each amplicon was digested with the appropriate restriction enzyme and purified using the GeneMATRIX DNA Purification Kit (EURx, Poland). In addition, the GFP fragment was defosphorylated using Calf Intestinal Alkaline Phosphatase (EURx) and ligated with the P2A sequence followed by PRKAA1 to obtain GFP-P2A-AMPKα1. Amplification of the entire construct was performed using attB primers, GFPattB forward (5`-GGGGGACAAAGTTTTGTACAAAGCAGGCTACCATGTGTGAGCAAG-GGCGA) and AMPKattB reverse (5`-GGGGACCACTTTTGTACAAGAAAG-CTGGGTTTATTGTGCAAGAATTTT), respectively. The purified product was directly recombined with the pDONR221 vector and then with the pLenti6/UbC-DEST expression vector using BP and LR clonase (Invitrogen, USA), respectively. The obtained clones were verified by restriction analysis and sequencing. A schematic of the GFP-P2A-AMPKα1 gene construct is shown in Figure S-3A.

***Production of viral vectors***

Lentiviral particles were generated accordingly to ViraPower protocol (Thermo Fisher Scientific) $9.5\times10^6$ HEK293T cells were seeded on T75 culture flask (DMEM high glucose, 10% FBS). Four hours after seeding, a transfection using calcium orthophosphate ($Ca_3(PO_4)_2$) was performed. The transfection mixture was prepared as follows: plasmid DNA ($1.5\times10^{12}$ copies of expression plasmid), 65 $\mu$l of 2.5 M $CaCl_2$, 650 $\mu$l of 2X BBS (BES buffered saline, pH 7.2) (both from Sigma-Aldrich, St. Louis, MO, USA) and 585 $\mu$l $H_2O$. The transfection mix was incubated for 20 min in room temperature, then transferred into fresh culture medium containing 25 $\mu$M chloroquine (Sigma-Aldrich). Medium containing lentiviral particles was harvested 48 and 72 h after transfection. Harvested medium was centrifuged (1750 g, 10 min, 4°C), filtered through 0.45 $\mu$m syringe filter and frozen at −80°C.

To obtain modified HTB-35 cell lines, cells were infected using generated lentiviral particles introducing the GFP-P2A-AMPK gene construct. Transduction was performed using MOI=5 (Multiplicity of infection) and 6 µg/ml polybrene (Sigma-Aldrich). Selection of HTB-35 HTB-35 GFP-P2A-AMPK modified cells was performed using the selection antibiotic blasticidin (Invivogen, USA) at a concentration of 6 µg/ml.

HTB-35 GFP-P2A-AMPK cell line was sorted using a FACSAria II cell sorter (Beckton-Dickinson) to enrich population of blasticidin resistant cell with cells of highest GFP expression (GFP expression is in line with AMPK expression due to the transgene design). 10% of the cells exhibiting the highest fluorescence has been collected. The percentage of green fluorescent cells in the modified cell lines was measured using an Attune NxT flow cytometer.

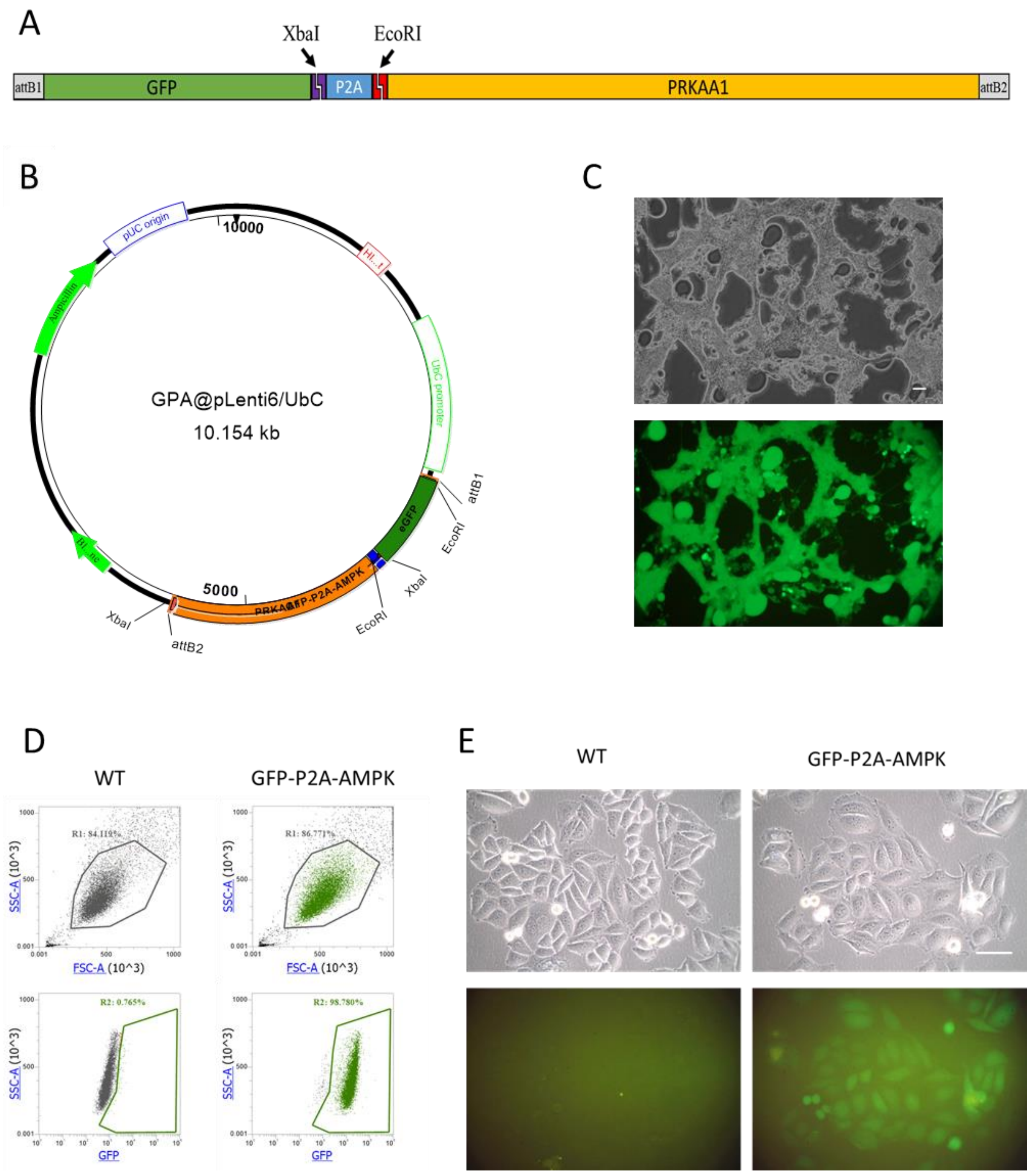


**Figure S- 3.** Generation of HTB-35 GFP-P2A-AMPK cell line. Design of GFP-P2A- PRKAA1 gene construct (A). Plasmid map of GFP-P2A-AMPK@pLenti/UBC expression plasmid (B). GFP-P2A-AMPK lentivirus production in HEK-293T cells (C) – visible syncytia due to lentivirus generation. Flow cytometry analysis of obtained HTB-35 GFP-P2A-AMPK cell line (D) and fluorescence of HTB-35 GFP-P2A-AMPK cells (E). White bars represent 100 μm.

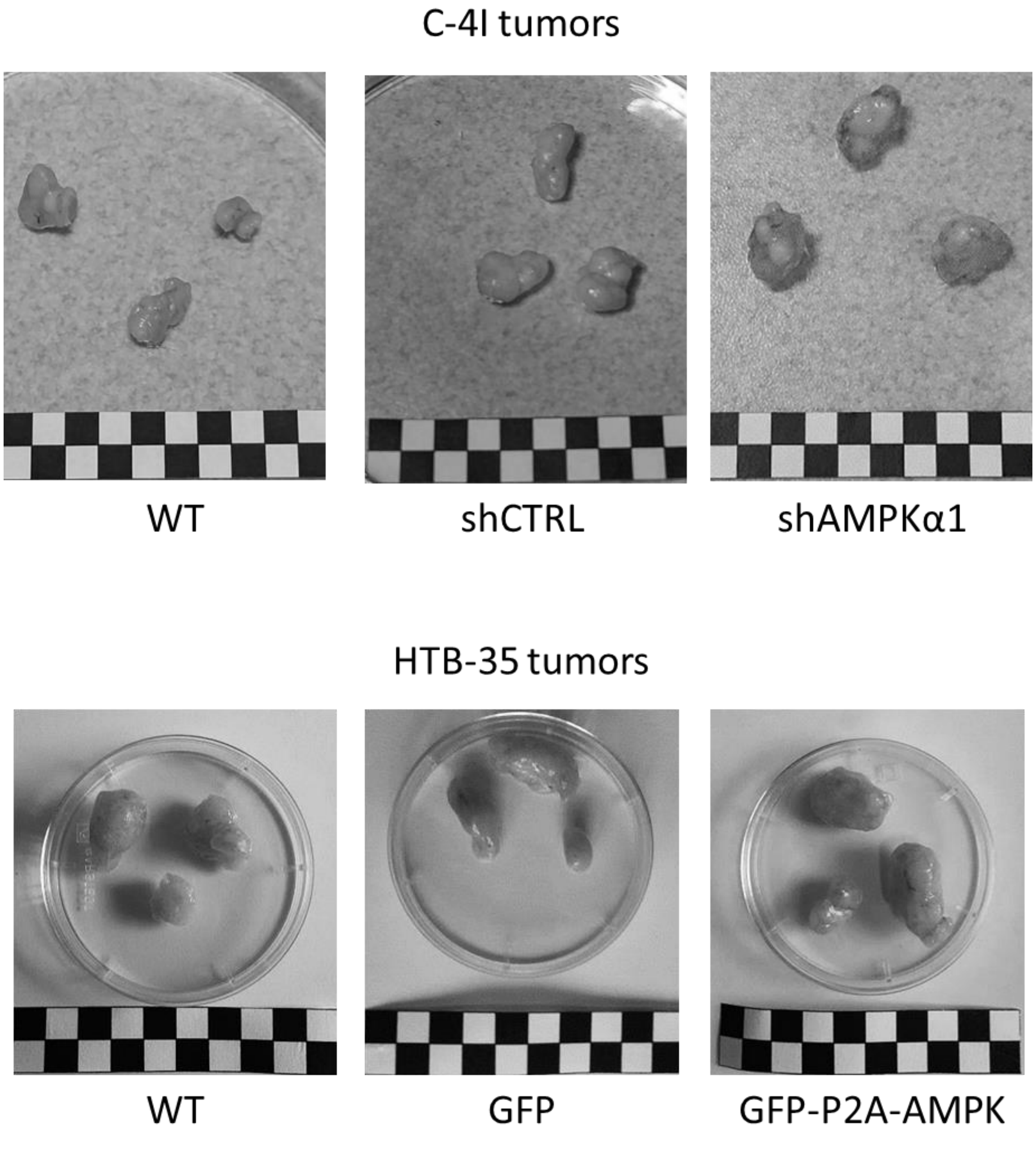


**Figure S- 4.** Tumors of C-4I and HTB-35 (WT and AMPK modified) cells extracted from mice. The sizes of black and white squares are 5x5 mm.
