## Supplementary material for "Impact of AMPK on cervical carcinoma progression and metastasis": Table 1

**Table 1. Taq-Man qRT-PCR probes list**

| **Gene name** | **Cat no.** | **Gene name** | **Cat no.** |
| --- | --- | --- | --- |
| PRKAA1 | Hs01562315_m1 | VIM | Hs00958111_m1 |
| PRKAA2 | Hs00178903_m1 | GFP | Mr03989638_mr |
| PRKAB1 | Hs00272166_m1 | DSP | Hs00950591_m1 |
| PRKAG1 | Hs01091629_g1 | CLDN1 | Hs00221623_m1 |
| SNAI1 | Hs00195591_m1 | OCLN | Hs00170162_m1 |
| SNAI2 | Hs00161904_m1 | DES | Hs00157258_m1 |
| CDH1 | Hs01023894_m1 | MMP-2 | Hs00234422_m1 |
| ZEB-1 | Hs00232783_m1 | ACTB | Hs99999903_m1 |
| MMP-1 | Hs00899658_m1 | MMP-13 | Hs00233992_m1 |
| mGAPDH | Mm99999915_g1 | hGAPDH | Hs02786624_g1 |
