## Supplementary material for "Impact of AMPK on cervical carcinoma progression and metastasis": Table 2

**Table 2. Antibodies utilized for protein detection**

| **Antibody target** | **Manufacturer** | **Cat no** |
| --- | --- | --- |
| AMPKα | Cell Signaling | #2793 |
| Phospho-AMPKα | Cell Signaling | #4188 |
| AMPKβ1 | Cell Signaling | #4178 |
| Phospo-AMPKβ1 | Cell Signaling | #4186 |
| GFP | Cell Signaling | #2955 |
| GAPDH | Cell Signaling | #2118 |
| goat anti-rabbit IgG-HRP | Santa Cruz Biotechnology | sc-2004 |
| goat anti-mouse IgG-HRP | Santa Cruz Biotechnology | sc-2005 |
